# Synergistic efficacy of tigecycline combined with apramycin against *tet*(X)-harboring *Acinetobacter spp.*

**DOI:** 10.1101/2023.07.20.549942

**Authors:** Juan Liu, Si-Lin Zheng, Jing-Jing Wu, Mei Zheng, Da-Tong Cai, Yan Zhang, Jian Sun, Ya-Hong Liu, Xiao-Ping Liao, Yang Yu

## Abstract

The emergence of the wide variety of novel tigecycline resistance *tet*(X) variants including *tet*(X3), *tet*(X4), *tet*(X5) and *tet*(X6) has posed a significant challenge to the clinical treatment of multidrug-resistant bacterial infections and represents a serious threat to global public health. The purpose of this study was to evaluate the synergism of tigecycline combined with other antibiotics as a means of overcoming *tet*(X)-mediated resistance in *Acinetobacter spp*. We found that the combinations of tigecycline with apramycin or amikacin exhibited synergistic activity against *tet*(X)-harboring *Acinetobacter spp*. with FICI values of 0.088 and 0.625, respectively. The MIC_TGC_ decreased >5-fold decrease in the presence of subinhibitory levels of apramycin. This combination was shown to be a therapeutically effective synergism using both *in vitro* and *in vivo* (mouse thigh infection model) assays and delayed the increase of MIC values over time. This study highlights the synergism of tigecycline in combination with apramycin which offers a viable therapeutic alternative for infections caused by *tet*(X)-harboring *Acinetobacter*.

## INTRODUCTION

Infections caused by *Acinetobacter spp.* first appeared in the 1960s and 1970s and have since become increasingly common worldwide (1). Initially, such infections were thought to be of low virulence potential or easily treatable with available therapies and any resulting mortality was often attributed to underlying patient conditions rather than the infection itself (2). However, the rapid emergence and global dissemination of *Acinetobacter spp.* as a major nosocomial pathogen is remarkable, particularly in intensive care units where it can cause illnesses such as pneumonia, tracheobronchitis, meningitis, endocarditis, peritonitis, skin and soft tissue infection and urinary tract and bloodstream infections (3)(4). In the United States, *Acinetobacter baumannii* is typically acquired through nosocomial infections and contributes to more than 99,000 deaths yearly (5). Consequently, there is an urgent need to develop new alternative strategies for treating these infections.

Currently, the options for combatting multidrug-resistant *Acinetobacter spp*. using antibiotics are limited. Tigecycline was approved for clinical use in 2005 and has since become an effective alternative for treating severe infections particularly those caused by extensively drug-resistant *Acinetobacter* strains (6)(7). Importantly, this last-resort antibiotic may become ineffective due to new tigecycline resistance *tet*(X) variants that include *tet*(X3), *tet*(X4), *tet*(X5) and *tet*(X6). Recent studies have highlighted the urgent need for new treatment options (8)(9).

Universally, treatment regimens for *Acinetobacter spp.* infections rely on colistin and tigecycline, either alone or in combination with carbapenems (10). Tragically, these last line agents for treating MDR-bacterial infections are increasingly challenged by the emergence and spread of antimicrobial resistance genes such as *tet*(X4)/(X5), *mcr* and *bla*_NDM_/*bla*_KPC_ (11)(12). In such situations, tigecycline-based regimens are alternative treatments against MDR bacteria (13)(14)(15) and it is crucial to explore effective drug combination to reduce the required tigecycline dosage.

Combination therapy is a method that involves the use of two or more active antibiotics in conjunction. This approach minimizes the dosage of toxic drugs and reduces the frequency of drug resistance, achieving more significant biochemical effects than monotherapy (16). For instance, a combination regimen of tigecycline and amikacin efficiently inhibited development of tigecycline resistance and successfully suppressed emergence of resistant populations (17). Tigecycline activity was also increased both *in vitro* and *in vivo* when used in combination with zidovudine against *E. coli* that carried *tet*(X) and *mcr-1* (18). Additionally, the use of apramycin in a mouse model of *A. baumanii* was effective when the AUC/MIC ratio was >50 and *C*_max_/MIC was 10 or higher (19).

There are only a limited number of studies exploring antibiotic synergy and its mechanisms Therefore, we sought to explore whether apramycin in combination with tigecycline can be effective against multidrug-resistant (MDR) *Acinetobacter spp.* strains harboring *tet*(X).

## RESULTS

### Bacterial Information

This study was conducted using 9 *Acinetobacter* isolates carrying different *tet*(X) variants to examine the effectiveness of combination therapy (Table 1). These included *Acinetobacter haemolyticus*, *Acinetobacter spp*. and *Acinetobacter beijerinckii* isolates carrying *tet*(X3), *Acinetobacter lwoffii* and *Acinetobacter indicus* isolates co-harboring *tet*(X3), *tet*(X6) and *bla*_NDM-1_ and three *A. indicus* isolates harboring *tet*(X3), *tet*(X6) / *bla*_NDM-3_ / *tet*(X4) and *tet*(X4) / *tet*(X5). All nine strains were resistant to tigecycline with MICs of 4-8 μg/mL but susceptible to colistin, polymyxin B, amikacin and apramycin. Four *bla*_NDM_-positive isolates exhibited resistance to meropenem.

**Table 1.**
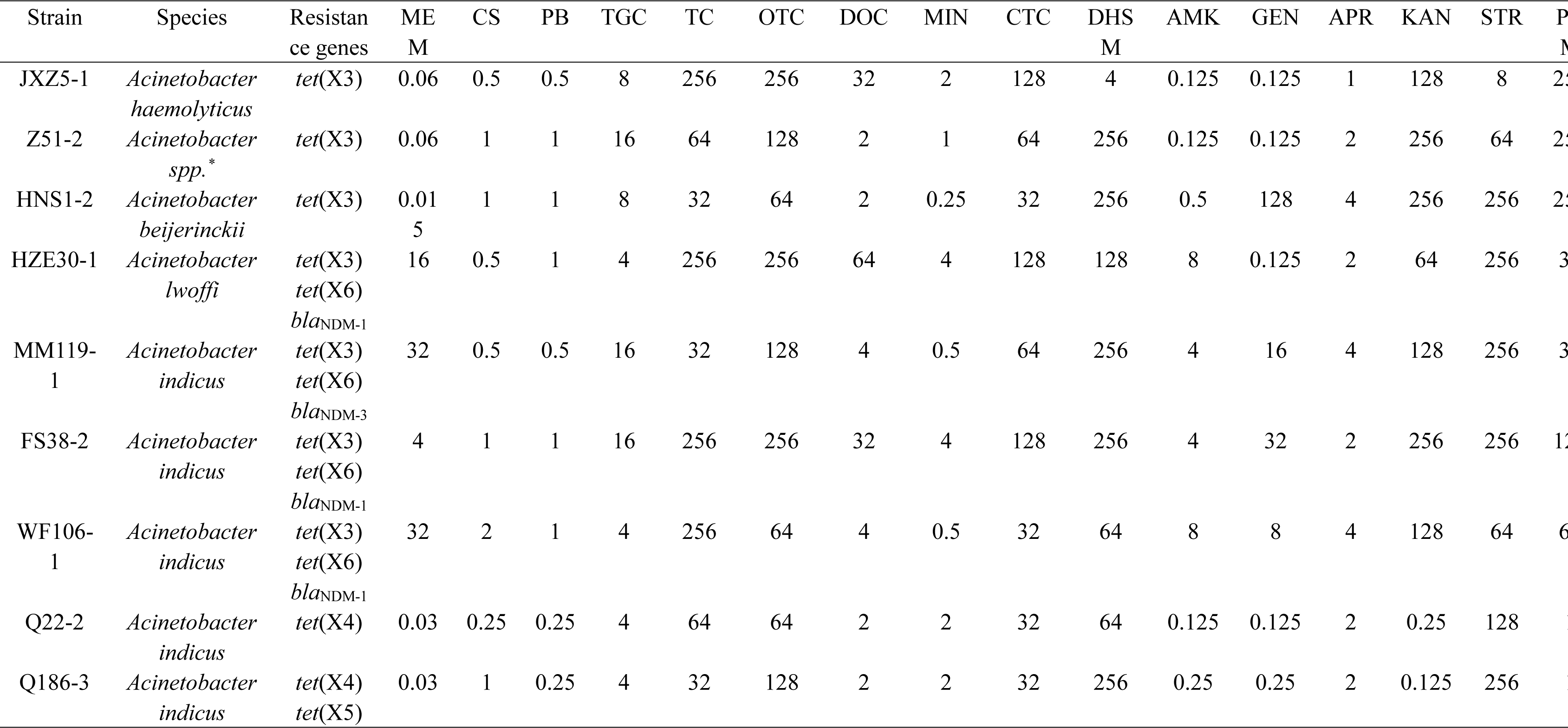
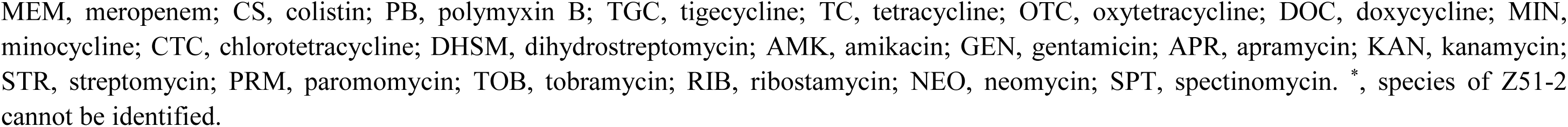
Molecular information and MICs of 9 *tet* (X)-harboring Aci*netobacter* strains.

| Strain | Species | Resistance genes | ME<br>M | CS | PB | TGC | TC | OTC | DOC | MIN | CTC | DHS<br>M | AMK | GEN | APR | KAN | STR | P<br>M |
| --- | --- | --- | --- | --- | --- | --- | --- | --- | --- | --- | --- | --- | --- | --- | --- | --- | --- | --- |
| JXZ5-1 | *Acinetobacter haemolyticus* | *tet*(X3) | 0.06 | 0.5 | 0.5 | 8 | 256 | 256 | 32 | 2 | 128 | 4 | 0.125 | 0.125 | 1 | 128 | 8 | 23 |
| Z51-2 | *Acinetobacter spp.\** | *tet*(X3) | 0.06 | 1 | 1 | 16 | 64 | 128 | 2 | 1 | 64 | 256 | 0.125 | 0.125 | 2 | 256 | 64 | 23 |
| HNS1-2 | *Acinetobacter beijerinckii* | *tet*(X3) | 0.01<br>5 | 1 | 1 | 8 | 32 | 64 | 2 | 0.25 | 32 | 256 | 0.5 | 128 | 4 | 256 | 256 | 23 |
| HZE30-1 | *Acinetobacter lwoffii* | *tet*(X3)<br>*tet*(X6)<br>*bla*<sub>NDM-1</sub> | 16 | 0.5 | 1 | 4 | 256 | 256 | 64 | 4 | 128 | 128 | 8 | 0.125 | 2 | 64 | 256 | 3 |
| MM119-1 | *Acinetobacter indicus* | *tet*(X3)<br>*tet*(X6)<br>*bla*<sub>NDM-3</sub> | 32 | 0.5 | 0.5 | 16 | 32 | 128 | 4 | 0.5 | 64 | 256 | 4 | 16 | 4 | 128 | 256 | 3 |
| FS38-2 | *Acinetobacter indicus* | *tet*(X3)<br>*tet*(X6)<br>*bla*<sub>NDM-1</sub> | 4 | 1 | 1 | 16 | 256 | 256 | 32 | 4 | 128 | 256 | 4 | 32 | 2 | 256 | 256 | 12 |
| WF106-1 | *Acinetobacter indicus* | *tet*(X3)<br>*tet*(X6)<br>*bla*<sub>NDM-1</sub> | 32 | 2 | 1 | 4 | 256 | 64 | 4 | 0.5 | 32 | 64 | 8 | 8 | 4 | 128 | 64 | 6 |
| Q22-2 | *Acinetobacter indicus* | *tet*(X4) | 0.03 | 0.25 | 0.25 | 4 | 64 | 64 | 2 | 2 | 32 | 64 | 0.125 | 0.125 | 2 | 0.25 | 128 | 1 |
| Q186-3 | *Acinetobacter indicus* | *tet*(X4)<br>*tet*(X5) | 0.03 | 1 | 0.25 | 4 | 32 | 128 | 2 | 2 | 32 | 256 | 0.25 | 0.25 | 2 | 0.125 | 256 | 1 |

MEM, meropenem; CS, colistin; PB, polymyxin B; TGC, tigecycline; TC, tetracycline; OTC, oxytetracycline; DOC, doxycycline; MIN, minocycline; CTC, chlorotetracycline; DHSM, dihydrostreptomycin; AMK, amikacin; GEN, gentamicin; APR, apramycin; KAN, kanamycin; STR, streptomycin; PRM, paromomycin; TOB, tobramycin; RIB, ribostamycin; NEO, neomycin; SPT, spectinomycin. \*, species of Z51-2 cannot be identified.

### *In vitro* checkerboard assays

Strain FS38-2 (*tet*(X3) *tet*(X6) *bla*NDM-1) was selected for screening the synergism of tigecycline combined with seven other antibiotics. The combination of tigecycline plus apramycin produced a synergistic effect (FICI = 0.088) while an additive effect (FICI = 0.625) was observed with amikacin. The other antibiotics with tigecycline produced interactive effects either indifferent or antagonistic (FICI ≥ 1). Additionally, further evaluation of tigecycline plus apramycin was repeated against all nine strains by checkerboard assays. This drug combination was synergistic against all 9 strains with an FICI _average_ of 0.590 (Figure 1b and Table S1).

**Figure 1.**
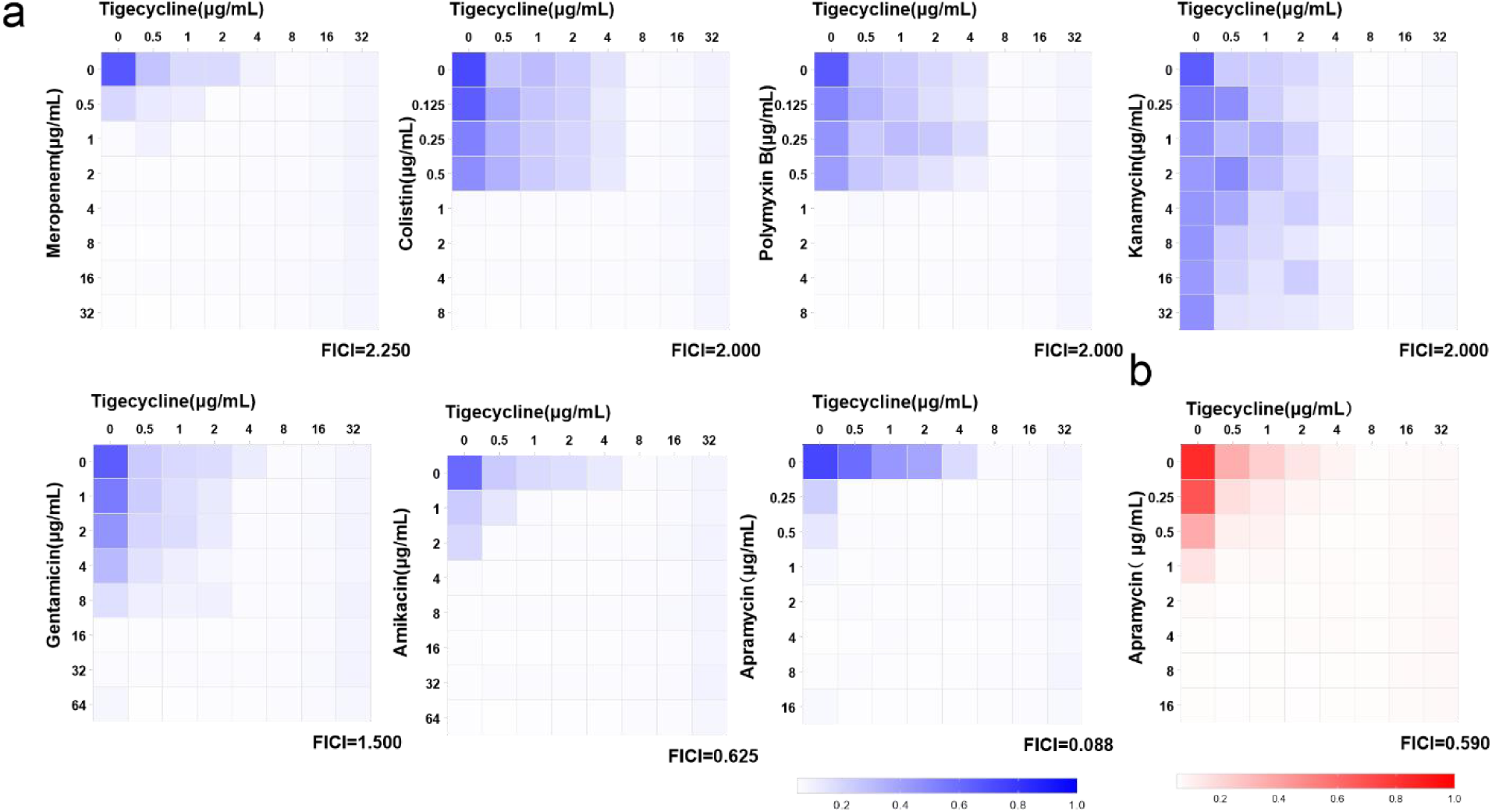
Heat-maps of checkerboard assays for tigecycline combined with the listed antibiotics. A. Strain FS38-2 was used in the preliminary screen b. Synergistic effect of tigecycline combining apramycin was further confirmed by checkerboard assays against all nine test strains (FICI_average_ =0.59, n=9 biological replicates).

### *In vitro* synergistic effectiveness

#### Combined MIC_TGC_ tests

The distribution of MIC_TGC_ against *Acinetobacter spp.* was evaluated when tigecycline was exposed alone or combined with 0.25, 0.5, 1, 2 μg/mL apramycin. As apramycin levels were increased, the MIC_TGC_ distribution shifted left on the curve towards lower MIC values of 0.25-0.50 μg/mL (Figure 2a). The magnitude of the MIC_TGC_ reduction when combined with apramycin was > 5-fold in the presence of subinhibitory levels of apramycin. Notably, tigecycline combined with 2 μg/mL apramycin resulted in > 20-fold reductions in MIC_TGC_ on average (n=9) and was significantly higher than with 0.25 μg/mL apramycin (*P* < 0.001) (Figure 2b).

**Figure 2.**
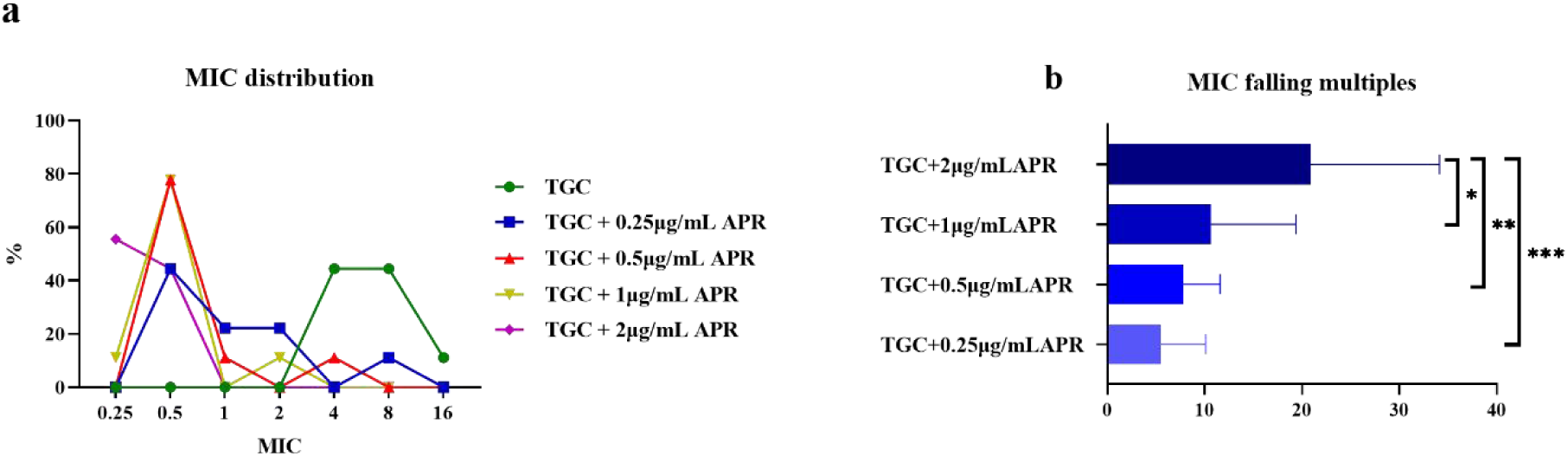
MIC changes of tigecycline against 9 *tet*(X)-harboring *Acinetobacter* strains when combined with apramycin at various concentrations. a. MIC distributions of tigecycline alone or combined with different doses of apramycin. b. MIC reductions of tigecycline in the presence of apramycin. \**P*<0.05, \*\**P*<0.01 and *\*\*\*P*<0.001. APR, apramycin; TGC, tigecycline; control, normal saline.

#### Time-kill and dose-response curves

We examined the bacteriostatic ability of APR/ TGC combinations using time-kill curves for all 9 *Actinobacter* strains. Tigecycline and apramycin levels of 16 and 4 μg/mL were completely growth inhibitory while lower concentrations displayed lesser effects as expected (Figure 3a and 3c). Apramycin at 0.5 μg/mL produced significant inhibitory activity in the presence of subinhibitory levels of TGC (Figure 3b). In contrast, a slight regrowth was detected using apramycin plus 0.5 μg/mL (Figures 3c and 3d).

**Figure 3.**
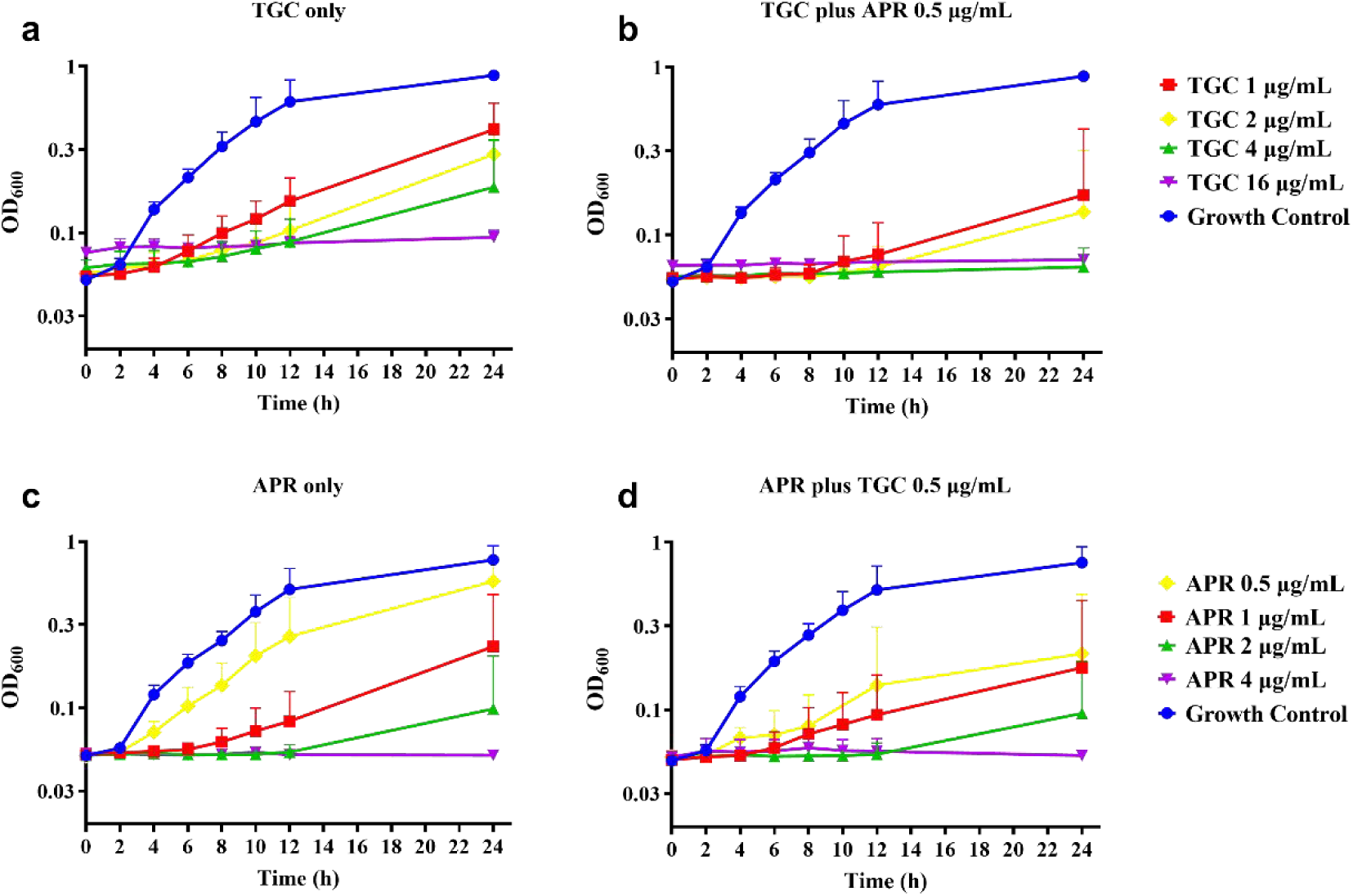
*In-vitro* time-killing curves of 9 *tet*(X)−harboring *Acinetobacter* strains with the indicated antibiotics. a. Tigecycline alone. b. Tigecycline combined with 0.5 μg/mL apramycin. c. Apramycin alone. d. Apramycin combined with 0.5 μg/mL tigecycline. APR, apramycin; TGC, tigecycline.

The concentration-effect relationship was fitted to a Hill-type equation in order to accurately determine the response to antibiotic challenge. Curves for tigecycline alone and tigecycline plus apramycin illustrated that greater inhibitory activity were obtained as the tigecycline concentration was increased (Figure 4a). Nonetheless, tigecycline doses > 0.5 μg/mL did not achieve better antimicrobial activity (Figure 4b) and this trend was consistent with the time-kill assay results. Importantly, compared with the single-drug groups, the IC_50_ of two-drug groupings (apramycin with 0.5 μg/mL tigecycline and tigecycline with 0.5 μg/mL apramycin) dropped to < 0.0001 μg/mL (Table 2).

**Figure 4.**
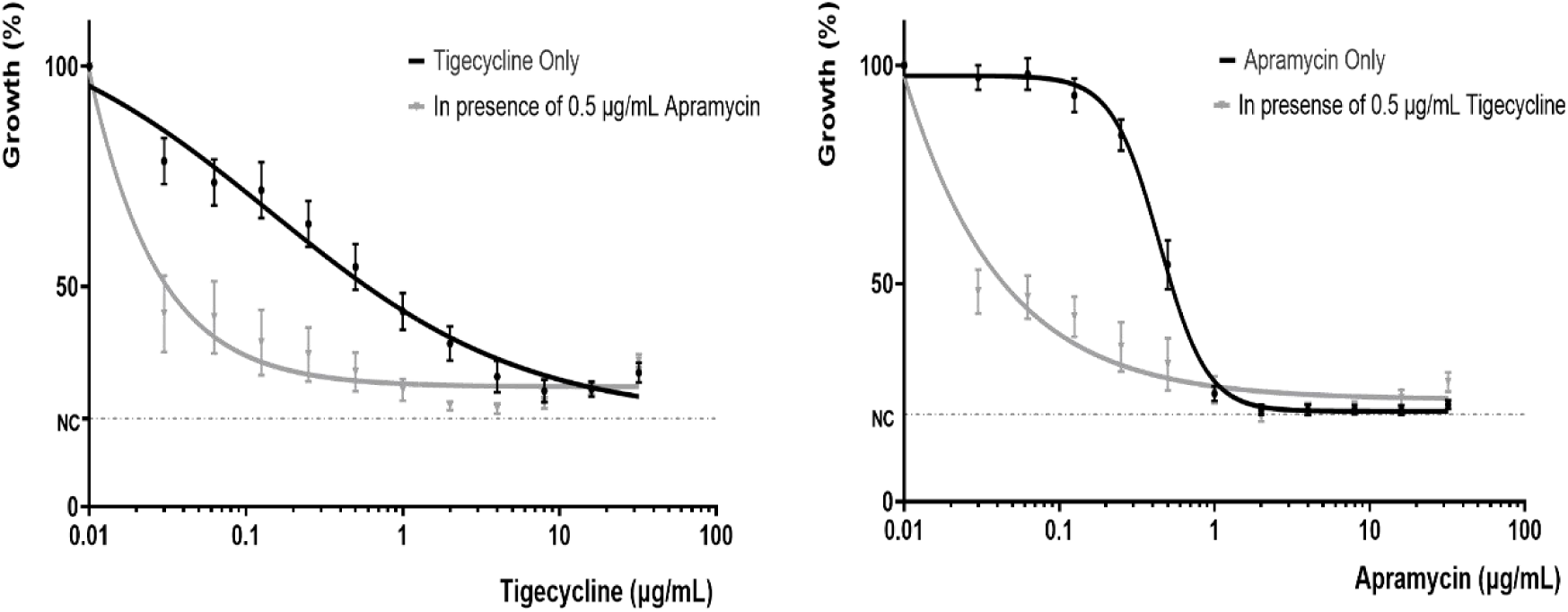
Dose response curves of different combinations of tigecycline and apramycin for *tet*(X)-harboring *Acinetobacter* strains. a. Dose response curve of tigecycline alone and in combination with apramycin (0.5 μg/mL); b. Dose response curve of apramycin alone and in combination with tigecycline (0.5 μg/mL). NC, negative control (minimum detection limit).

**Figure 5.** Combination therapy of tigecycline plus apramycin in a mouse thigh model. See Figure 2 for abbreviations.

**Table 2.**
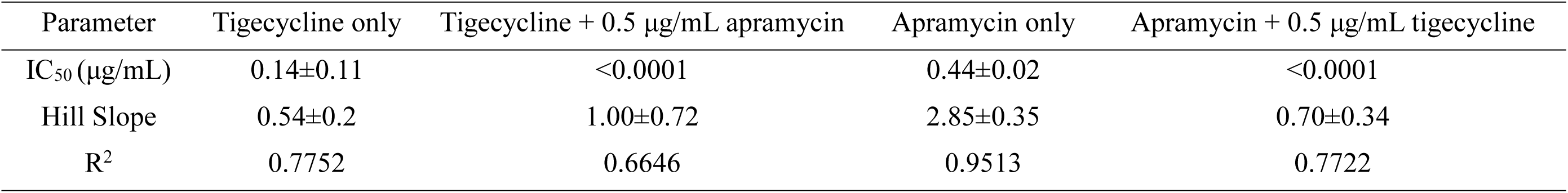
Dose-response parameters of different combinations of tigecycline and apramycin against 9 *tet*(X)-harboring *Acinetobacter*.

| Parameter | Tigecycline only | Tigecycline + 0.5 µg/mL apramycin | Apramycin only | Apramycin + 0.5 µg/mL tigecycline |
| --- | --- | --- | --- | --- |
| IC<sub>50</sub> (µg/mL) | 0.14±0.11 | <0.0001 | 0.44±0.02 | <0.0001 |
| Hill Slope | 0.54±0.2 | 1.00±0.72 | 2.85±0.35 | 0.70±0.34 |
| R<sup>2</sup> | 0.7752 | 0.6646 | 0.9513 | 0.7722 |

### *In vivo* synergistic efficacy

We next examined whether this synergism would also be apparent *in vivo* using a neutropenic mouse thigh model. The bacterial density in the control groups were maintained at 10^6^-10^7^ CFU/g after 24 h treatment with control saline. In contrast, bacterial growth was reduced to 10^2^10^3^ CFU/g using combination therapy against all four test strains (*P* < 0.001). Tigecycline or apramycin monotherapies inhibited growth of all test strains except HNS1-2. The combination therapies displayed robust killing activity demonstrating synergism of tigecycline plus apramycin (*P <* 0.01). However, we found significant differences in bacterial growth density only with tigecycline alone and in combination against strains Q186-3 and HZE30-1 (*P* < 0.01). Considerable antibacterial effects were observed with tigecycline and apramycin combined although only slight combined activity was seen with strain Q186-3 *tet*(X4) / *tet*(X5) (Figure 5). These data indicated that combination therapy groups generated significant bacterial killing (P < 0.05).

### Prevention of high-level tigecycline-resistant mutants

One important aspect of combination therapy development is to ensure that high-level resistance is not generated. When we exposed our test strains to tigecycline alone, the MIC_TGC_ for most of the strains showed rapidly fluctuating and rising trends. This was especially apparent for strains MM119-1 and Z51-2 whose MIC_TGC_ reached 64 μg/mL (8 × initial MIC_TGC_) on days 3 and 4. The addition of 0.5 μg/mL apramycin generated MIC_TGC_ values centered at 4 μg/mL. This indicated the strains were losing sensitivity to tigecycline with continuous exposure to the drug combination. Notably, during the 14 days of passaging only strain JXZ5-1 was completely inhibited by the combined drugs (Figure 6).

**Figure 6.**
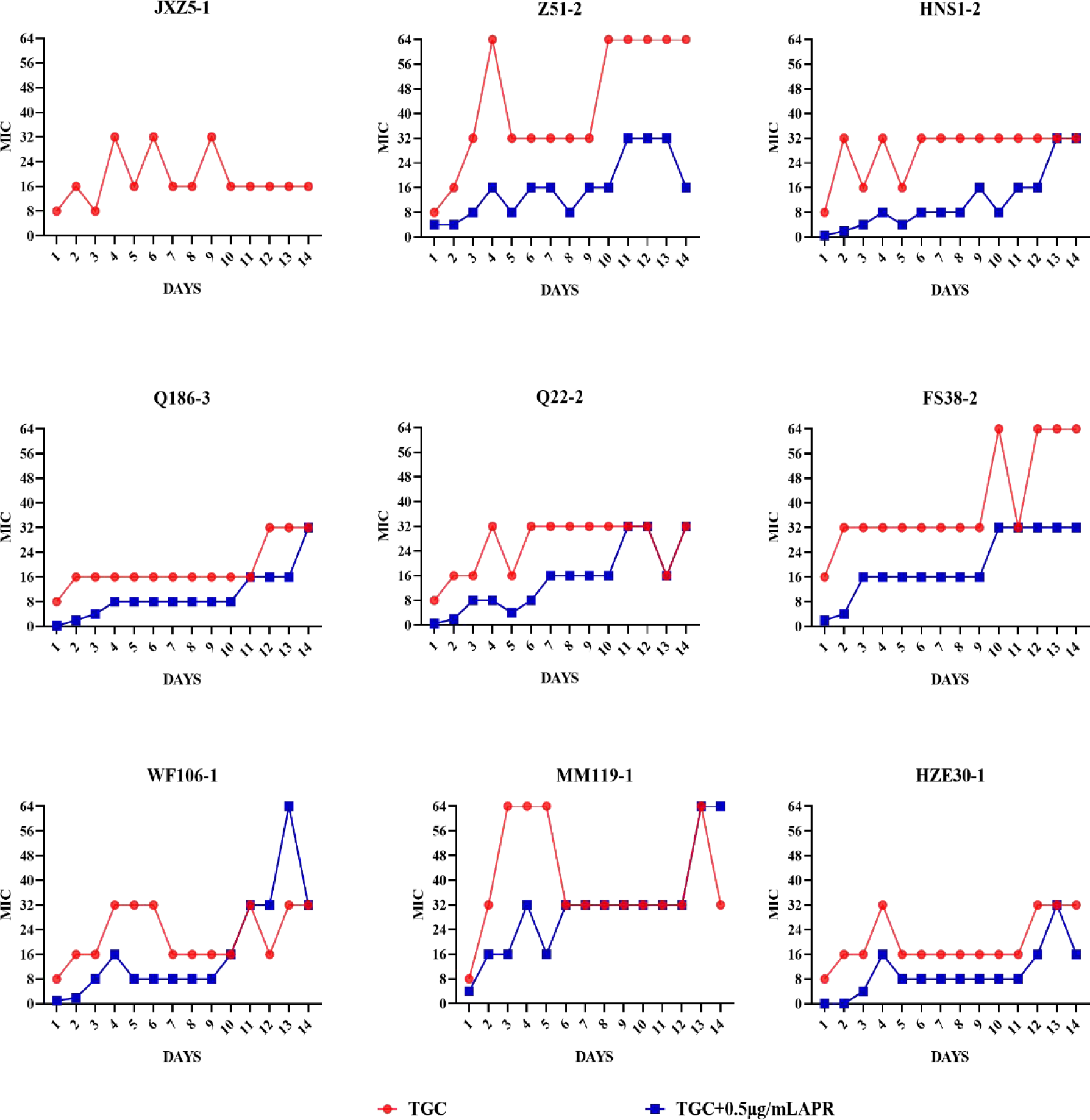
MIC_TGC_ changes of *tet*(X)-harboring *Acinetobacter* strains under drug selective pressure. Red, tigecycline alone; Blue, tigecycline plus 0.5 μg/mL apramycin. Strain JX5-1 could not be passed continuously under the combined drug pressure and bacterial growth was completely inhibited by the APR/TGC combination.

## DISCUSSION

The majority of the *tet*(X)−harboring *Acinetobacter* strains are associated with livestock raising concerns that the tigecycline-resistance *tet*(X) genes could be transmitted to humans through opportunistic pathogens like *Acinetobacter* (10)(20). f (15). Additionally, infections caused by non-fermentative Gram-negative bacilli have been treated successfully with aminoglycosides based on tigecycline treatment results (21). Our results prove that these combinations are also effective against *Acinetobacter* strains possessing *tet*(X) (Figure 1b and Table S1).

Combination therapy has become a commonly employed approach for the treatment of severe infections caused by *Acinetobacter spp*. (22). In our study, when partnered with apramycin, certain strains reverted to the tigecycline sensitive phenotype especially with the addition of 2 μg/mL apramycin and resulted in a 20-fold lower MIC_TGC_ (Figure 2). Since dose response models in preclinical trials can be predictors of drug interactions (23), we used time-kill curves and dose response models for screening. We found that as the drug concentration was increased, the antibacterial effect of combined drug group exceeded that of the single drug groups. This accentuated bacteriostatic effect of tigecycline became increasingly apparent in the apramycin combinations (Figures 3 and 4, Table 2). Moreover, a recent report demonstrated the potential of combining colistin/amikacin and tigecycline to achieve synergistic effects against *Acinetobacter* strains further support our findings (24).

While the initial response of patients to a combination therapy often appears promising, the emergence of resistance to continuous therapy is not uncommon (25). In our group of *tet*(X)-carrying isolates, one was significantly inhibited or killed by tigecycline plus 0.5 μg/mL apramycin during the 14 days of passaging and only 22.2% of isolates possessed a high tigecycline MIC (64 μg/mL) with continuous exposure of the combination (Figure 6). This indicated that apramycin may delay the MIC increase of tigecycline.

Further confirmation <u>o</u>f the antimicrobial activity of the combination therapy was carried out using an *in vivo* mouse model. Prior to treatment, four strains with different resistance *tet*(X) genotypes colonized the mouse thigh as expected. The drug combinations demonstrated significant antibacterial efficacy when compared to the controls (Figure 5). Previous reports have indicated that elevated MICs of either apramycin or tigecycline would dampen their clinical efficacy when utilized as a monotherapy option (26)(27)(28). Indeed, we found that strain HNS1-2 (MIC_TGC_ 8 μg/mL, MIC_APR_ 4 μg/mL) that carried *tet*(X3), displayed a highly significant synergistic response to this combination *in vivo* although tigecycline monotherapy failed to produce positive outcomes. In contrast, 3/4 strains remained resistant to tigecycline. Fortunately, the combination of tigecycline and apramycin exerted strong synergistic effects *in vivo*. Differences of 3-5 log CFU/g were achieved between control groups and the combined treatment groups and differences of 1-5 log CFU/g were observed between monotherapies and the combination therapies. Supporting our study, the synergistic activity of two antibiotics may be related to co-resistance genes and the MIC of tigecycline (15)(29). Aminoglycosides inhibit protein synthesis by binding to the 16S rRNA and disrupt the integrity of bacterial cell membranes that may facilitate tigecycline passive accumulation (30). During combination therapy, tigecycline dominated the bacterial responses and might lead to synergistic bactericidal effects by enhancing the aminoglycoside effect and inhibiting bacterial adaptive responses (31). Further research is needed to determine the exact mechanisms for the success of this combination therapy.

In conclusion, the results of our study demonstrate the ability of apramycin and amikacin to provide synergistic activity with tigecycline. We provide evidence for the effectiveness of tigecycline combined with apramycin against *tet*(X)-harboring *Acinetobacter spp.* infections that also mitigated increases in MIC values. This study provides insights into a reassessment of tigecycline as a therapeutic option to treat *Acinetobacter* infections.

## METHODS AND MATERIALS

### Bacterial profiling and antimicrobial susceptibility

A total of 9 *Acinetobacter* isolates carrying different *tet*(X) variants were stored in a 30% glycerin broth and preserved at −80℃. The bacterial species were identified by matrix-assisted laser desorption/ionization time of flight mass spectrometry (MALDI-TOF, MS). The presence of *tet*(X) and co-harboring with *bla_NDM_* were identified by polymerase chain reaction (PCR) amplification and sequencing as previously reported (11). Antimicrobial susceptibility tests of tigecycline, meropenem, colistin, polymyxin B and the aminoglycosides amikacin, gentamicin, apramycin, kanamycin were performed using the broth microdilution method in Mueller-Hinton medium according to EUCAST guidelines. Reference strain ATCC 25922 served as quality control.

### Checkerboard assays and fractional inhibitory concentration index (FICI)

The checkerboard assay was used to test the joint antibacterial activity of tigecycline combined with seven antibiotics (see above). The reasons for these tests were as follows: (i) these drugs remained active against multidrug-resistant (MDR) *Acinetobacter* isolates (32)(33); (ii) some of them, like meropenem and colistin are still considered last resort treatments of infections caused by MDR Gram-negative bacteria (13)(3); (iii) few studies have evaluated the synergistic potential of tigecycline combinations against *Acinetobacter spp.* (15)(34).

In brief, isolate FS38-2 (MIC_TGC_=16 μg/mL and co-harboring *tet*(X3), *tet*(X6) and *bla*_NDM-1_ genes) was randomly selected for screening the synergic partners combining with tigecycline for the other seven antibiotics. The checkerboard assays of the potential drug combinations were performed against all nine isolates to further confirm the synergism. Bacterial cultures of 5×10^5^ CFU/mL were exposed to serial 2-fold dilutions of antibiotics with concentrations ranging from 1/8× to 8×MIC in 96-well plates and incubated for 18 h at 37°C. The FICI index was analyzed by the following equation:

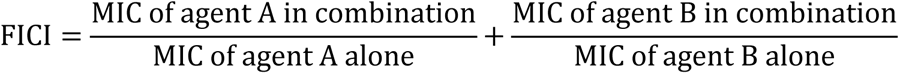

where FICI ≤ 0.5 represents “synergism”; 0.5 < FICI≤ 1 means “addition”; “indifferent” was defined when 1< FICI ≤ 2; and FICI> 2 denotes “antagonism”(35).

### Time-kill assays

The antibacterial time-kill assay was used to assess the results of the antibiotic monotherapies and combination against 9 *tet*(X)-harboring *Acinetobacter* isolates. This study was composed of four groups: Groups A and B, only tigecycline, tigecycline plus 0.5 μg/mL apramycin (Final tigecycline levels; 0, 0.03, 0.06, 0.12, 0.25, 0.5, 1, 2, 4, 8, 16 and 32 μg/mL); Group C and D, only apramycin, apramycin combined with 0.5 μg/mL tigecycline (Final apramycin levels, the same as tigecycline Groups A and B). An initial inoculum of ∼10^5^ CFU/mL incubated with different drug concentrations at 37°C and the OD_600_ nm readings to trace bacterial proliferation were made at 0, 2, 4, 6, 8, 10, 12 and 24 h using an Ensight Multimode Plate Reader (PerkinElmer, Waltham, MA, USA).

### Dose–response assays

As described previously (36),OD values at hour 8 was selected to calculate the correlation between antimicrobial effects and dosage of drugs with formula using GraphPad Prism Version 8. The OD values for bacterial growth in the absence of antibiotics was used as the normalization standard. Results are shown as the mean value of all 9 isolates at a 95% CI. The dose-response correlation of tigecycline or apramycin was calculated with the equation:

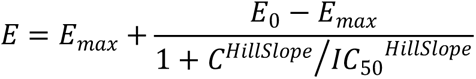

where *E* is the bacterial growth estimated as the OD_600_ normalized by that of control; *E_max_* is the maximal antimicrobial efficacy; *E_0_* is the bacterial growth in the absence of any antibiotics; *C* is tigecycline or apramycin concentration; *IC_50_* is the tigecycline or apramycin concentration demonstrating the 50% of *E_max_*; Hillslope describes the curve gradient.

### *In vivo* therapeutic verification

The neutropenic mouse thigh model was employed for testing the *in vivo* synergistic efficacy of tigecycline plus apramycin. Specific-pathogen-free (SPF) female ICR mice weighing 25±2 g at six-weeks-old (Hunan Silaikejingda Lab Animal, Hunan, China) were rendered temporarily neutropenic by immunosuppression with cyclophosphamide (Yuanye Biotechnology, Shanghai, China) at 150 mg/kg on the first four days and 100 mg/kg at the fifth day by intraperitoneal injection before bacterial inoculation. In respect of the animal ethics and saving cost, four different *tet*(X)-carrying *Acinetobacter* isolates were selected for *in vivo* experiments. The four isolates were *A. lwoffi* HZE30-1 and *A. indicus* WF106-1 (both harboring *tet*(X3), *tet*(X6) and *bla*_NDM-1_), *Acinetobacter* HNS1-2 carrying *tet*(X3) and *A. indicus* Q186-3 carrying *tet*(X4) and *tet*(X5). The mid-log bacterial cultures were appropriately diluted with normal saline and the neutropenic mice (neutrophils ≤ 100/mm^3^) were then intramuscularly inoculated with 100 µL of bacterial suspension (10^7^ CFU /mL) into each posterior thigh muscle. After a 1 h, the control normal saline Group I or antibiotics were administered in the following manner: single-drug groups received only tigecycline (Group II), apramycin (Group III); and combined groups received tigecycline combination with apramycin (Group IV). Four mice were tested in each group. Dosing regimens were 5 mg/kg for tigecycline administrated subcutaneously, 20 mg/kg apramycin subcutaneously and the injection volume was 100 µL for all drugs. The tigecycline doses were selected to mimic the pharmacokinetic profiles of recommended human clinical doses (37)(38). In the case of apramycin, the doses were selected based on previous reports of pharmacodynamic effects on different MIC and preliminary experimental results (19)(39). Following 24 h of treatment, mice were sacrificed and thigh homogenates were sampled for bacterial burden quantification.

### Prevention of high-level tigecycline-resistant mutants

To further investigate whether antibiotic combinations prevent the occurrence of high-level tigecycline-resistant mutants, nine strains were challenged with prolonged and repeated exposure to tigecycline plus apramycin. Approximately 5×10^5^ CFU/mL of *tet*(X)-carrying *Acinetobacter* isolates cells were challenged by tigecycline only and tigecycline plus 0.5 μg/mL apramycin. After incubating at 37°C for 18 hours, cultures from each MIC well and the last well containing bacterial growth were mixed and challenged with the antibiotic groups described above. This protocol was repeated continuously for 14 days.

## ETHICS APPROVAL

The *in vivo* mouse studies were approved by the Animal Research Committee of South China Agricultural University and the Guangdong Association for Science and Technology [ID: SCXK (Guangdong) 2018-0002]. All experiments have followed the guidelines of Guangdong Laboratory Animal Welfare and Ethics and the Institutional Animal Care and Use Committee of the South China Agricultural University [2021F138].

## TRANSPARENCY DECLARATIONS

None to declare.

## Supplementary material

**Table S1.** MIC and FICI values of TGC/AP against *tet*(X)-harboring *Acinetobacter* strains.

| Strain | Resistance gene | TGC | APR | FICI |
| --- | --- | --- | --- | --- |
| JXZ5-1 | *tet(X3)* | 8 | 1 | 0.3125 |
| Z51-2 | *tet(X3)* | 16 | 2 | 2 |
| HNS1-2 | *tet(X3)* | 8 | 4 | 0.3125 |
| HZE30-1 | *tet(X3) tet(X6)*<br>*blaNDM-1* | 4 | 2 | 0.25 |
| MM119-1 | *tet(X3) tet(X6)*<br>*blaNDM-3* | 16 | 4 | 0.375 |
| FS38-2 | *tet(X3) tet(X6)*<br>*blaNDM-1* | 16 | 2 | 0.75 |
| WF106-1 | *tet(X3) tet(X6)*<br>*blaNDM-1* | 4 | 4 | 0.375 |
| Q22-2 | *tet(X4)* | 4 | 2 | 0.75 |
| Q186-3 | *tet(X4)*<br>*tet(X5)* | 4 | 2 | 0.1875 |

